## Supplementary material for "Characterization of GEXP15 as a potential regulator of Protein Phosphatase 1 and partner of ribosomal complex in *Plasmodium falciparum*": BioRxiv Supplementary material.pdf

### 1 Supplementary Figures

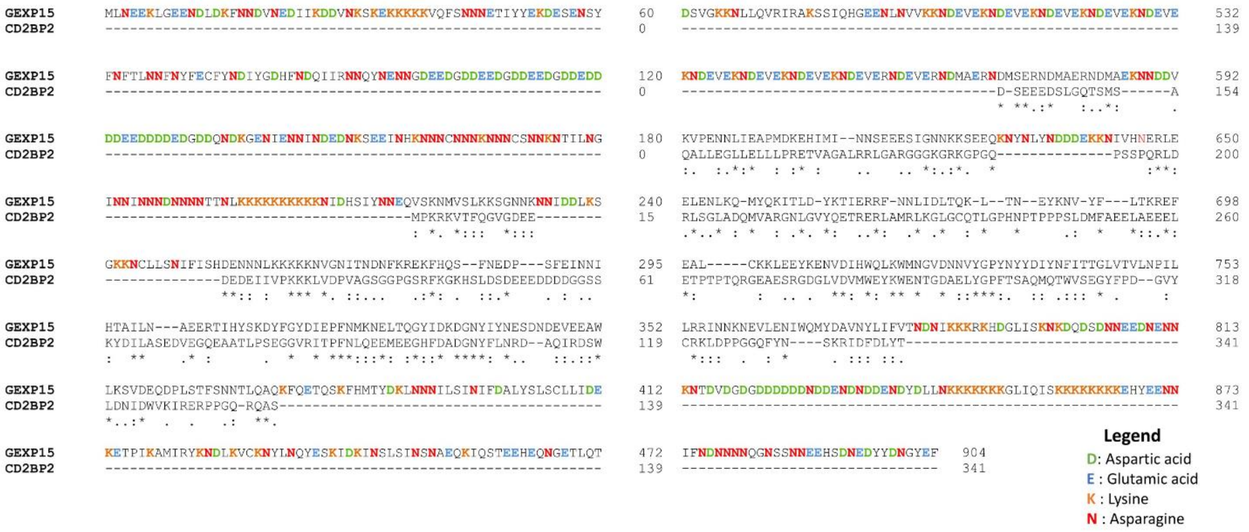


**Supplementary Figure S1. The protein sequence alignment of PfGEXP15 and HsCD2BP2.** The alignment was performed using ClustalW. Colored amino acids represent low complexity regions.

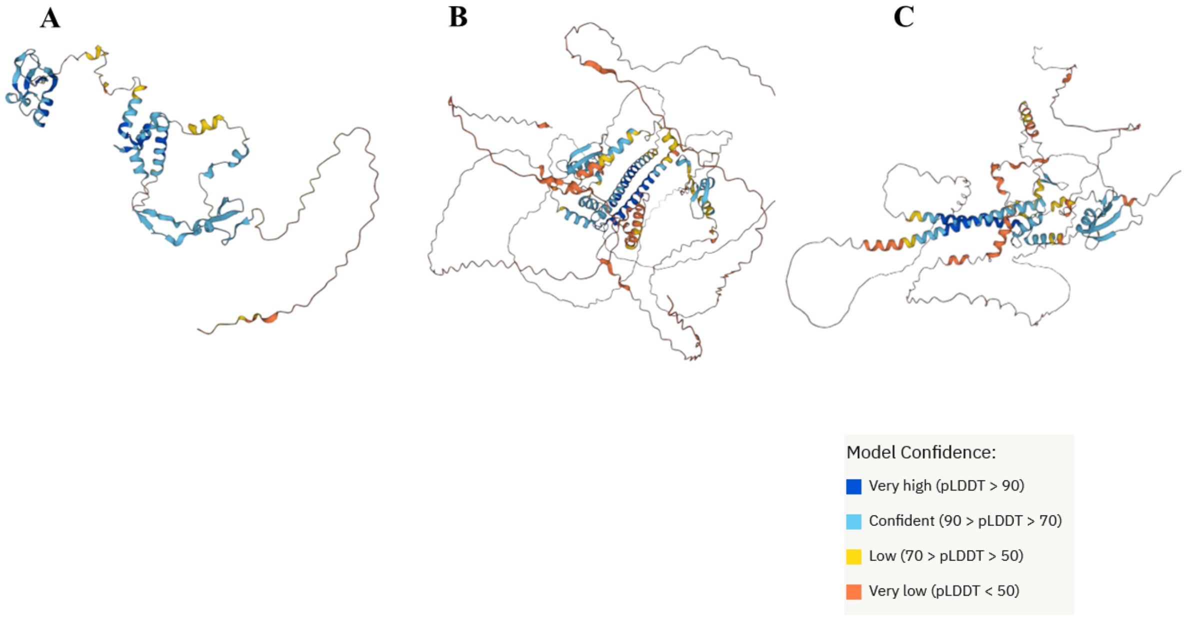


**Supplementary Figure S2. The 3D structure prediction of HsCD2BP2, PfGEXP15 and PbGEXP15.** The models of HsCD2BP2 (A), PfGEXP15 (B) and PbGEXP15 (C) were generated by AlphaFold.

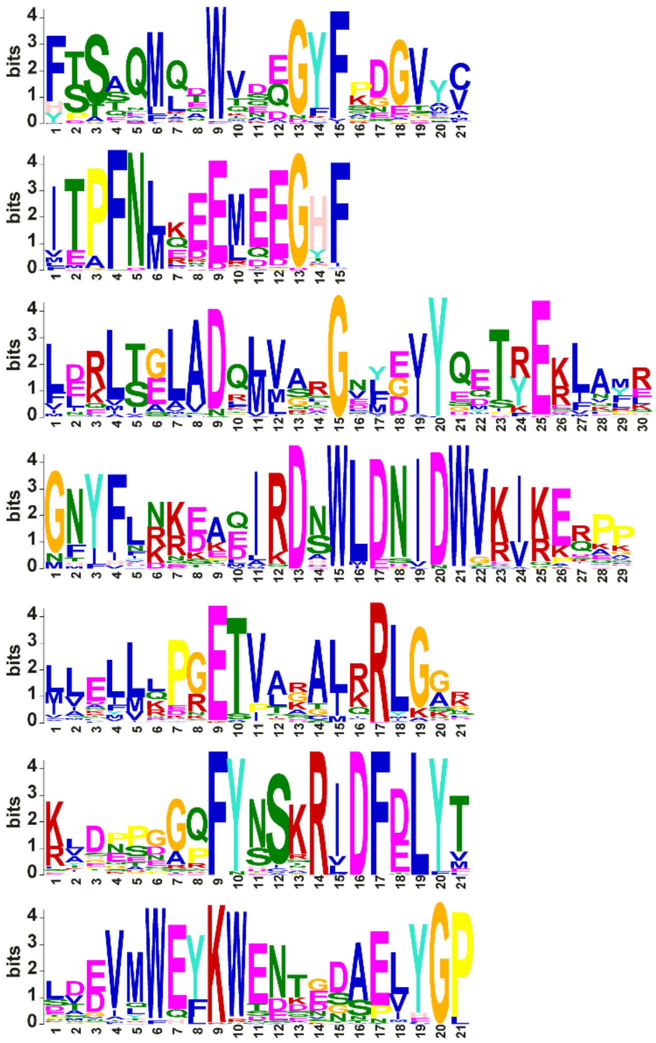

| Motif | E-value | Sites | Width |
|---|---|---|---|
| 1 | 7.6e-878 | 75 | 21 |
| 2 | 2.3e-778 | 79 | 15 |
| 3 | 2.6e-1000 | 70 | 30 |
| 4 | 3.2e-975 | 54 | 29 |
| 5 | 2.2e-751 | 80 | 21 |
| 6 | 1.5e-666 | 55 | 21 |
| 7 | 3.1e-653 | 56 | 21 |

**Supplementary Figure S3. MEME motif search of CD2BP2 homologs in eukaryotes.** The 7 most significant sequence logos identified by MEME are represented, as well as their respective E-value, number of sites and width across the protein sequences. The height and size of the letters represent the amino acid frequency.

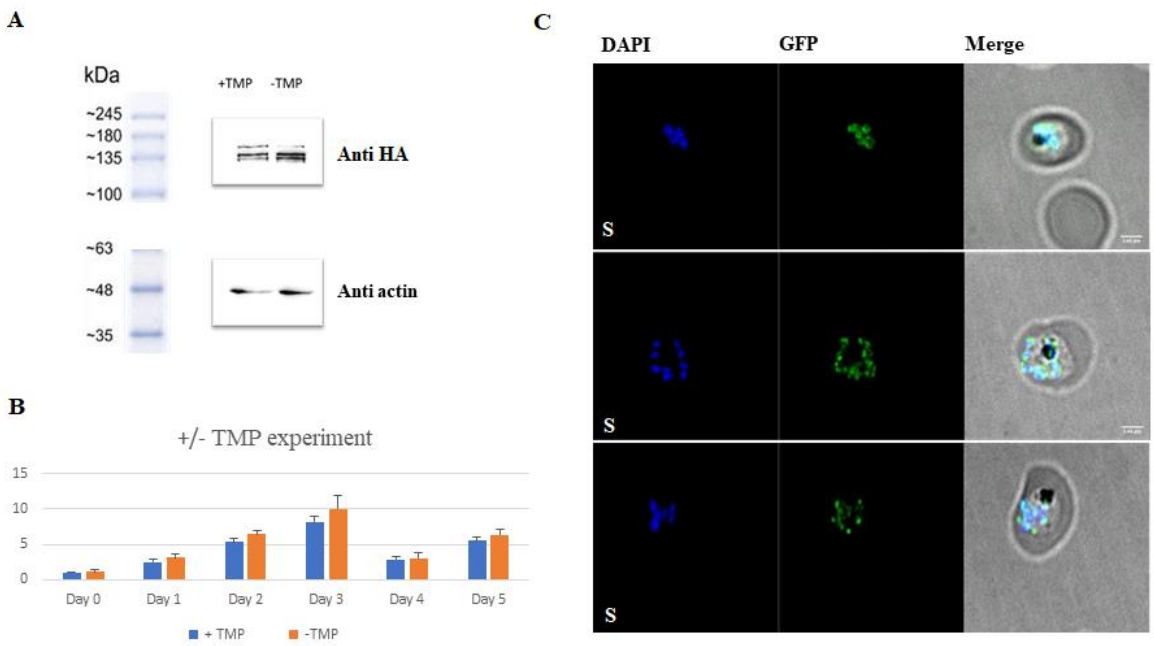


**Supplementary Figure S4. TMP removal does not affect parasite growth and PfGEXP15 localization A.** Western Blot analysis representing the total protein extract from a highly enriched cultures of transgenic iKd PfGEXP15 late trophozoites with TMP in lane 1 and without TMP for 12 days (corresponding to around 7 cycles) in lane 2. They were revealed with mAb anti-HA. In the lower panel, anti-actin was used as a positive loading control. 40 million parasites were used in each lane. **B.** Parasitemia of iKd PfGEXP15 line was measured with and without TMP cultures. The results are shown as the mean parasitemia ± SD. (n=4). **C.** Confocal laser scanning microscopy showing GFP expressing parasites in transfected cultures without TMP. Parasite's nuclei were stained with DAPI and transgenic parasites are expressing PfGEXP15-GFP-DDD-HA. Merged images showed the protein colocalization.

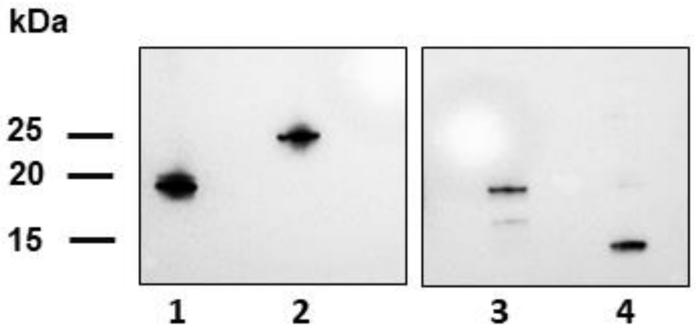


**Supplementary Figure S5.** Western blot analysis representing RVxF motif (lane 1), tetR protein (lane 2), UD (lane 3) and GYF domain (lane 4) recombinant proteins eluted from nickel beads and detected using anti-His antibodies. The figure was set up from the same western blot at different exposures.

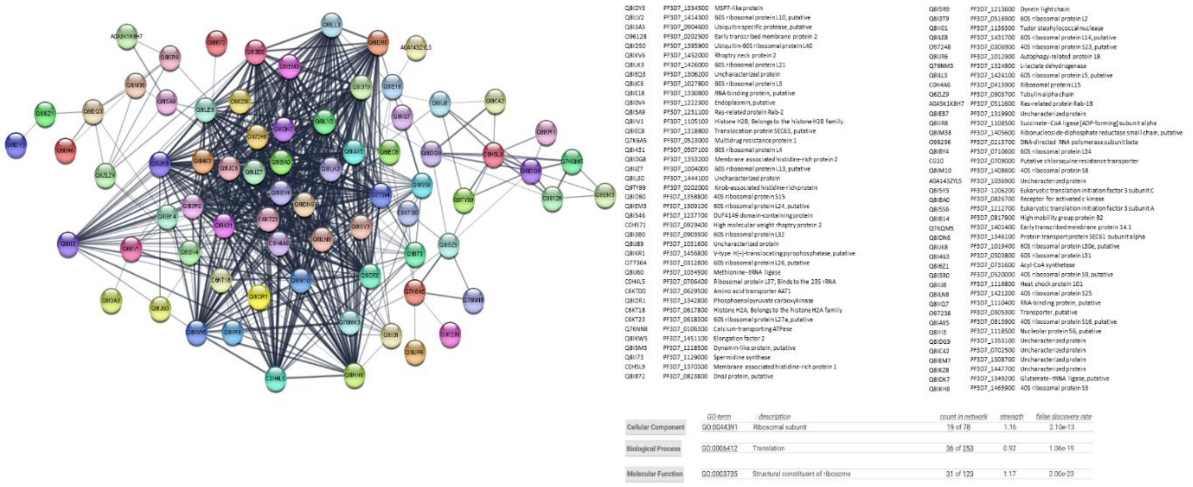

**Supplementary Figure S6. GYF-domain containing protein pulldown analysis. A.** STRING network visualization of GYF-interacting proteins using Cytoscape software **B.** List of GYF interacting partners, as well as the processes they may be involved.

### 2 Supplementary Datasheets

Supplementary Datasheet S1.xlsx

Supplementary Datasheet S2.xlsx

### 3 Supplementary Tables

Supplementary Table S1.xlsx

Supplementary Table S2.xlsx

Supplementary Table S3.xlsx

Supplementary Table S4.xlsx

Supplementary Table S5.xlsx
